# Semiconducting bacterial biofilm based on graphene-MoS2 template and component dependent gating behavior

**DOI:** 10.1101/2020.09.13.295360

**Authors:** Sanhita Ray, Arpita Das, Anjan Kr Dasgupta

**Affiliations:** Department of Biochemistry, University of Calcutta, Kolkata; Department of Electrical Engineering, Jack Baskin School of Engineering, University of California, Santa Cruz

## Abstract

In this paper, we report for the first time, the synthesis of a semiconducting biofilm. Photosynthetic bacterial biofilm has been used to weave together MoS2 nanosheets into an adherent film grown on interdigitated electrodes. Liquid-phase exfoliation of bulk MoS2 powder was used to obtain MoS2 nanosheets. A synchronous-fluorescence scan revealed the presence of two emission maxima at 682nm and 715nm for the MoS2 suspension. Such maxima with bandgap energy 1.82 and 1.73 eV corresponded to the single and double layer of MoS2. The presence of such single and multi-layered structures was confirmed by Raman spectroscopy, FTIR studies, and electron microscopy. The current-voltage (I-V) studies of such a bio-nano hybrid revealed the emergence of the gated nature of the current flow. This Schottky diode like behavior, reported earlier for Graphene-biofilm junctions, is also observed in this case. Gating voltage depended on the composition of the biofilm. The semiconductor biofilms, when studied using electrochemical impedance spectroscopy, revealed characteristic Nyquist and Bode plots, suggesting special circuit-equivalence for each film. While Mos2 was marked with stability with respect to variations in RMS voltage and bias voltage, the graphene biofilm was unique by the absence of any Warburg element.

## Introduction

The emergence of novel electrical properties in the interface of a 2D material like graphene and a biologically evolved 2D material like biofilm (1) has been recently reported by our group. Neuron like dynamic electrical behavior of biofilms (2) has made the 2D bio-nano hybrid more notable and enabling material.

The effort to replace 2D material like graphene by *MoS*2 has been continuing from few years. The 2D character of *MoS*2 and its role as a highly efficient fluorescence quencher has been emphasized (3) from the sensor context. The present paper for the first time explores the characters of *MoS*2 embedded biofilms which can exhibit more versatile behavior than *MoS*2 loaded with smart biomolecules. One such strange emergence is the presence of gated current flow like the neuron firing.

Biofilms are communities of bacterial cells arranged in fractal architecture, providing mechanical integrity and protection to individual cells. They consist of cells and extracellular materials (ECM), which is mainly comprised of proteins and polysaccharides(4). Rhodobacter capsulatus is a gram-negative, photosynthetic, microaerophilic purple non-sulfur bacteria that is fortunately non-pathogenic in nature.

We reported its time-dependent fluorescence at 613 nm (5). The photon is received by antenna pigments which then gets transported to the reaction center (6). The energy is stored by electron transfer from bacteriochlorophyll to an electron acceptor molecule through several subunits and cofactors. As another attribute of biofilms, we reported the specificity of a template (e.g. defects when studying graphene biofilm interaction) on photosynthetic metabolism (7).

Molybdenum disulphide (*MoS*_2_) is a transition metal dichalcogenide (TMD) bearing the formula *MX*_2_. The TMDs have unsaturated d-orbitals and defects at the edge sites which impart semiconducting or metallic behavior. Monolayer TMDs can be synthesized either by top-down(exfoliation from bulk) or bottom-up(synthesis using precursors) approach on a specific substrate (8).

*MoS*_2_ monolayers are stacked via Van der Waal’s force of interaction within bulk multilayered structures. *MoS*2 monolayer has a direct bandgap of 1.8 eV and exhibits semi-conducting behavior. This is in contrast to graphene which has indirect bandgap and has so far been the focus material for synthesizing hybrid biofilms. This bandgap makes *MoS*2 an ideal sensor material. The number of monolayers can be deduced from Raman spectra by determining the distance between the two fingerprint peaks at 384 *cm*_1_ and 405 *cm*_1_ (9). Exfoliation of *MoS*2 into monolayer or few layers imparts *MoS*2 with unique mechanical, chemical, optical, and electronic properties (10).

A number of biosensors have been fabricated based on *MoS*_2_, for sensitive qualitative or quantitative assay. Biomolecules play a vital role in metabolism. A 3D-*MoS*2/rGO/Au composite derived sensor was designed over a glassy carbon electrode and exhibited electrocatalytic activity for ascorbic acid(AA), dopamine (DA), and uric acid (UA) detection (11).

Another group of researchers proposed gold sensors modified with monolayer *MoS*2. Such sensors were reported to detect DNA molecules selectively. Since different photoluminescence (PL) properties were detected towards single and double-stranded DNA by monolayer *MoS*2 nanosheet, this property was utilized to design the biosensor depicting DNA fragment detection. This sensor can also be used to distinguish target DNA from mismatch DNA through the photoluminescence spectra of *MoS*2. Loan et al designed a graphene/*MoS*2 composite biosensor for label-free and selective detection of DNA hybridization through measuring the change of PL peak intensity.

While these specific sensing effects are important in the present paper we propose to use the *MoS*2 biofilm hybrid as a platform for material characterization, which in turn can be scaled up to a smart biosensing device.

## Materials and methods

### Liquid Phase Exfoliation

We have used a top-down approach to get the nanosheet structures using liquid-phase exfoliation. An NMP:water mixture in a ratio of 1:100 was made and the materials were added in this mixture. A water bath was prepared at 70 and 55C, while the mixtures were exfoliated extensively. The NMP and water react to produce hydroperoxides which aid in exfoliation of the materials. However, NMP solely would behave like a poor solvent. 55mMOl of water was used since this mixture will be incorporated directly into a bacterial culture and a high concentration of NMP would kill bacterial cells.

### Bacterial growth

The Rhodobacter culture was mentained using RCB mediamalic acid(4gm/l),Ammonium chloride(1gm/l), Calcium chloride(CaCl2.2H2o)(75mg/l), Nicotinic acid(1mg/l), Na-EDTA(20mg/l), Yeast Extract(0.3 g/l), MgCl2(120mg/l), KH2PO4(684.7 mg/ml), K2HPO4(865.7 mg/l) (pH-6.8) at 27C with constant light exposure (see S1, in the supplimentary to note how growth was monitored by absorbance or fluoresence)

### Scanning Electron Microscopy

In order to fix the samples for SEM imaging, the nanocomposite was grown over a glass surface. For fixation, the sample was washed in MilliQ and then kept immersed in 2.5% Glutaraldehyde for 15 minutes. After fixation, the sample was washed with MilliQ water for 4-5 times. Then the samples were treated with 1% Osmium Tetraoxide(OsO4). The sample then went through a series of diluted alcohol solutions (50%,70%,90%,100%) to remove water. SEM images were obtained using a scanning electron microscope (Model ZEISS EVO-MA 10). Images were obtained after platinum sputtering over the fixed samples in order to make the sample conductive and resistant to burning up.

### Transmission electron microscopy

TEM image of the *MoS*2 was obtained by drop-casting 5ul of exfoliated *MoS*2 sample over the TEM grid and the degree of exfoliation (Number of layers) was drying in a lightbox. EDAX was also performed to determine atomic ratios 80 within the material.

### Raman spectroscopy

Silicon chips were cut from a P-type silicon wafer. These chips then went through a series of washing procedures. These chips were washed thoroughly with hydrofluoric acid to remove the silicon oxide layer of the chips or else the nanoparticles will not be able to adhere to the surface of the chip. These chips were then washed with ethanol, 2-propanol and acetone sequentially to remove any dirt and adhered organic substances, hydrofluoric acid, or other contaminations. The silicon chips were placed on a Petri dish and 5-10 microlitre of diluted samples were then added over the shiny surface of the silicon chip. The sample was dried and characterized using Raman spectroscopy(Lab RAM HR Jovin Yvon) using a laser at 488nm.

### Fluorescence and photoluminescence

Bacterial fluorescence at 613 nm was measured by UV lamp excitation at 395 nm. Exfoliated *MoS*2 suspension generated a lot of scattering hence its PL was characterized by using a synchronous fluorescence method. In this method, excitation and emission are shifted simultaneously keeping the wavelength difference the same.Figure S2 in the supplementary shows how MoS2 leads to quenching of fluoresence in some region while amplification in some other region of the spectrum.

### Bio-fabrication on bioelectronic devices

Interdigitated electrodes (IDE) were obtained from NanoSPR LLC and screen-printed electrodes (SCE) were obtained from Dropsens (DRP-550). After the washing procedure alcohol was removed from the surface by letting it dry inside laminar airflow, in a sterile environment. Rhodobacter was grown over the surface of the electrodes by incubating it in 11ml of media (with 4 ml inoculum), containing 10mg/ml of *MoS*2, in a 15 ml falcon tube. After an incubation period of 5 days, electrodes were taken out from the suspension and washed, prior to performing electrical measurements.

### Preparations for impedance spectroscopy

We employed an interdigitated electrode (IDE) from DropSense which had an inter-finger gap of 10 microns, fingers of 20 microns, and a total width of 1mm, a dimension of 28mm × 5mm × 0.4mm with a 150nm thick layer of gold on 5nm Chromium underlayer. The frequency-dependent electrical measurements were taken on a Keithley SCS-4200 using Tektronix Parameter Analyzer software.

Impedance spectroscopy (IS) is a tool for investigating the electrical and electrochemical properties of materials and systems(12). Essentially the spectroscopic approach consists of the rendering of a Nyquist plot (a plot of the imaginary part of impedance Z plotted against the corresponding real part). The Nyquist plot in turn can be approximated to an equivalent circuit. Interestingly, the soft material in question may show complex frequency dependence of the real and imaginary part of impedance, and therefore the circuit may be more complex than expected. A typical example of such a circuit may be illustrated by

In the circuit we may explain the *Z*”(*ω*) and *Z*’(*ω*) by a circuit like *R*0 − *p*(*R*1 − *C*1) − *p*(*R*2 − *Z*_*W*_, *C*2).In this representation the element *Z*_*W*_ takes account of the diffusion, transport, migration between boundaries of the complex semiconductor-biofilm milieu. We may approximately call this a Randles circuit (13), a popular equivalent circuit in electrochemistry and electrochemical impedance spectroscopy that takes account of semi-infinite diffusion-controlled reaction occurring in a planar electrode. We can briefly express *Z*_*W*_ = *A*_*W*_*/*(*jω*)^0.5^, where *A*_*W*_ is Warburg coefficient, *ω*angular frequency).

## Results

### Electron Microscopy

In TEM imaging, 2-layered *MoS*2 was observed. it was approximately 0.44 *µm* and 0.67 *µm* across (see figure 1).In SEM imaging, the *MoS*2 imbedded *Rhodobacter* biofilm forms a bridge between the two fingers of of the interdigited electrode. The *MoS*2 platelets are held tightly onto the surface of the electrodes by the Rhodobacter biofilm cells.

**Figure 1:**
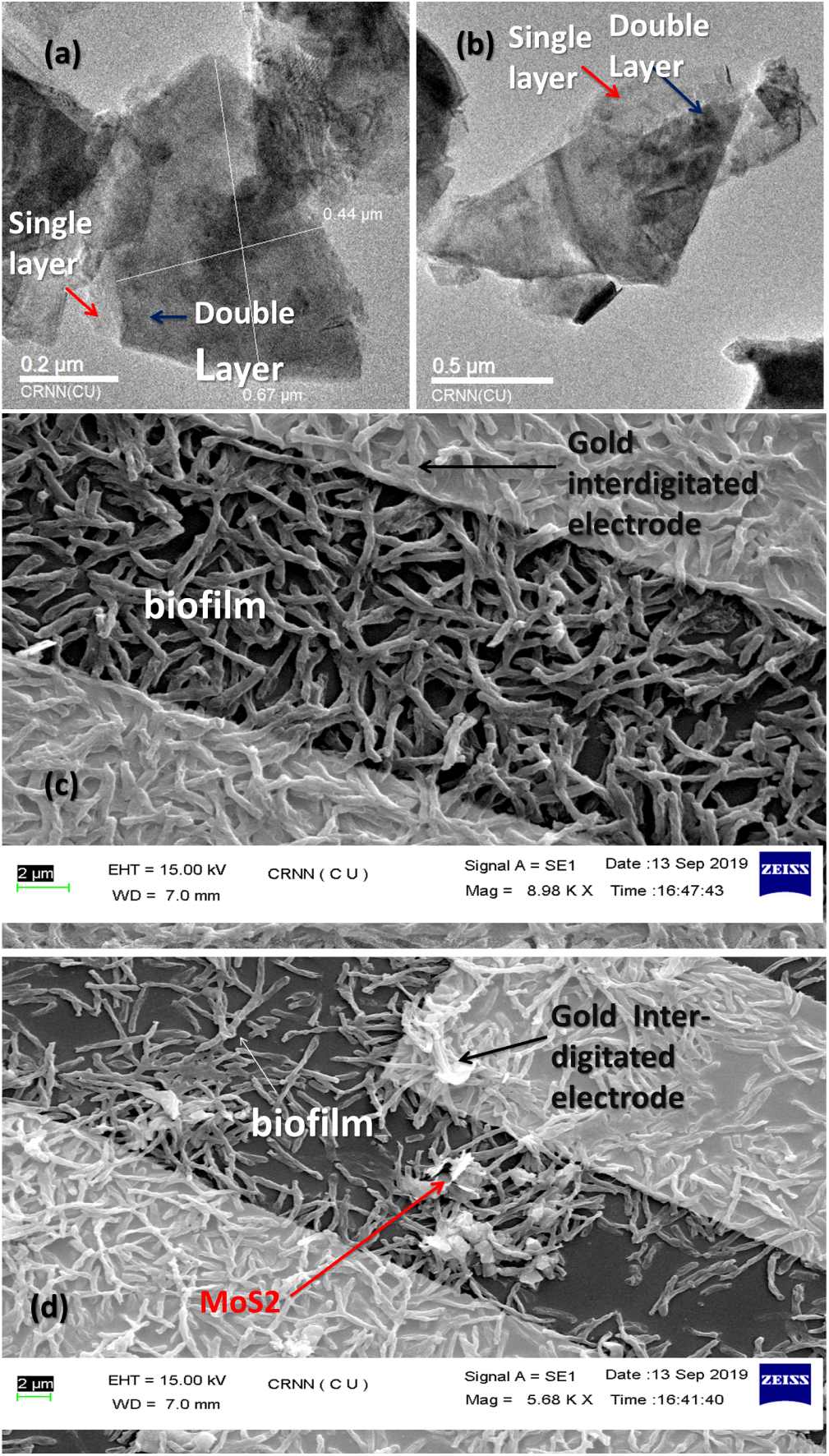
TEM image of liquid exfoliated *MoS*2 (a) and (b). SEM image of bacterial growth over interdigited fingers of IDE electrodes (c). SEM image of *MoS*2 integrated biofilm over IDE electrodes

### Fourier-transform infrared (FTIR) spectroscopy

FTIR was done to determine the chemical bonds in *MoS*2 and their interaction with NMP-Water mixture(figure 2a). Peaks at 2853*cm*^−1^ and 2925*cm*^−1^ signifies the presence of mono-layer *MoS*2 which is not found in bulk *MoS*2 samples according to (14).Weak peak of *MoS*2 was observed at 438*cm*^−1^ contributong to Mo-S vibration (9). Other peak at 618*cm*^−1^ which signifies multi-layered *MoS*2(14).The peak at 1628*cm*^−1^ marks the presence of *MoS*2 water bond (15,16). A very strong and broad signal at 3440*cm*^−1^ was observed which signifies the OH stretching. This could be due to the presence of minute amount of water in the exfoliated *MoS*2 sample (17).A strong prominant peak at 1382*cm*^−1^ is due to stretching vibration of methyl group of NMP and a peak at 1121*cm*^−1^ signifies in-plane bending of C-H bond (9).

**Figure 2:**
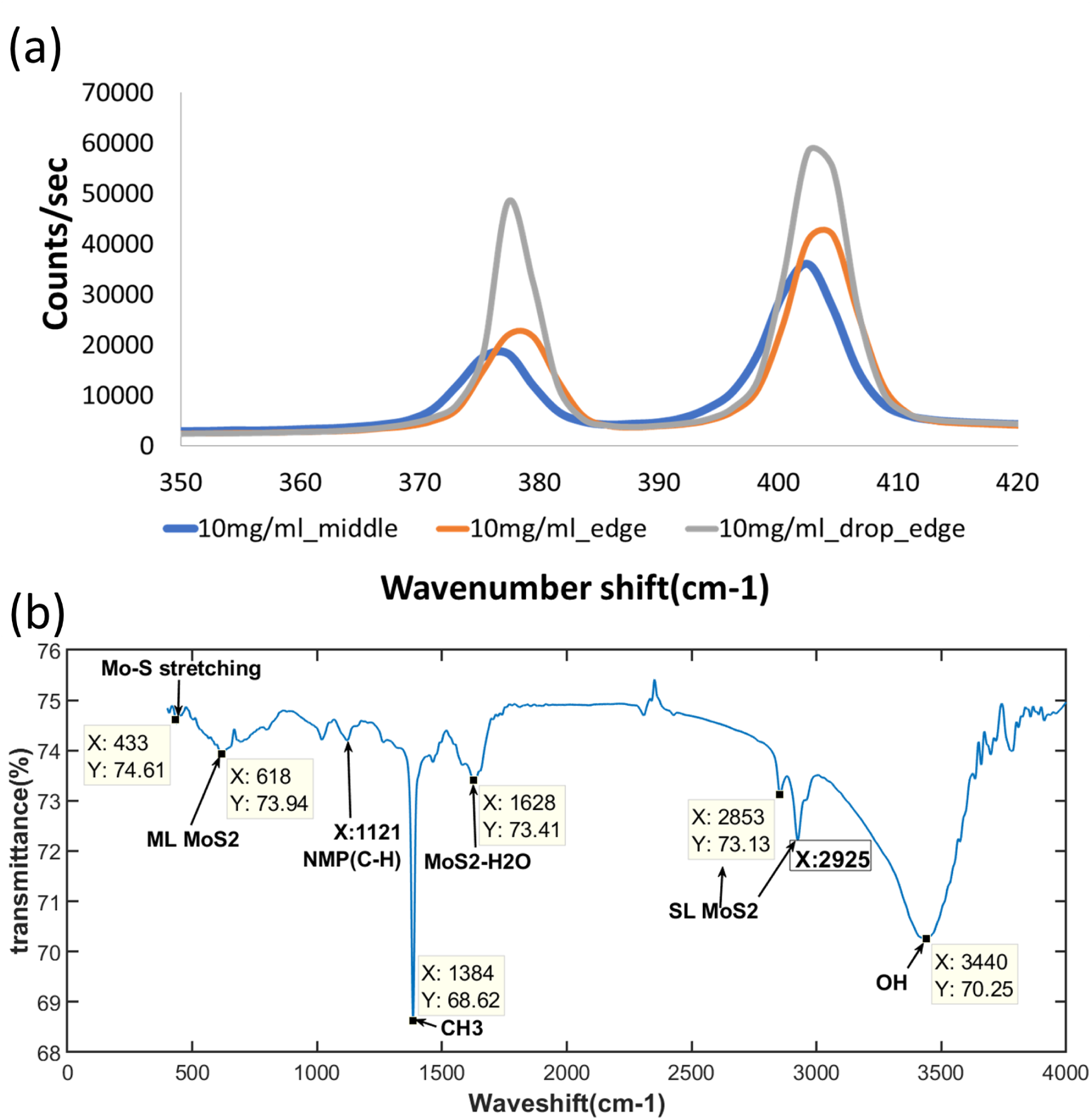
(a)Raman spectroscopy of exfoliated *MoS*2 taken at different positions of silicon waffer after dropcasting(b)FTIR spectroscopy exfoliated *MoS*2 (ML=Multi layer)(SL= single layered *MoS*2

**Figure 3:**
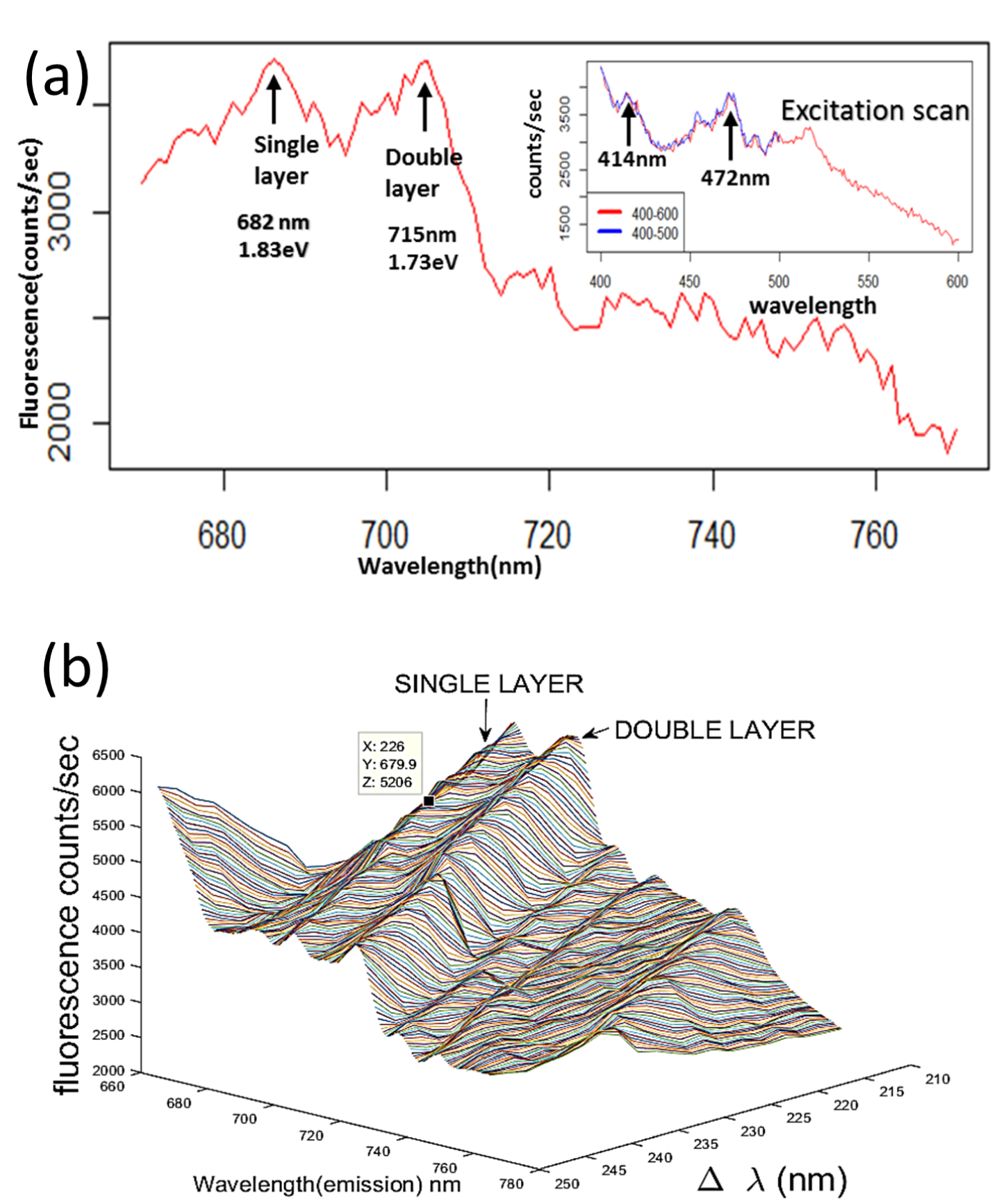
Synchronous fluorescence spectrum of liquid-phase exfoliated *MoS*2 drop cast on silicon wafer

### Raman Spectroscopy

The peak difference (in wavenumbers)in the fingerprint region of Raman spectrum,namely,370-410 *cm*^−1^, is found to be 25.09 *cm*^−1^ (Figure 2b).The reported difference (18,19) for a mono-layer is however 19.8cm^−1^. The difference may be accounted for by the presence of monolayer as well as multilayers of *MoS*2. The presence of such mixed layers is also revealed in electron microscopic studies (figure 1)) and fluorescence spectroscopic studies (see the next section).

### Fluoresence Spectroscopy

*Rhodobacter* shows fluorescence peak at 613 nm. *MoS*_2_ upon forming hybrid with Rhodobacter quenched the 613 nm fluorescence. A synchronous scan of *MoS*_2_ suspension in NMP: water mixture and observed it’s maximum fluorescence peak at 685nm. Upon doing a synchronous scan (438-538:670-770nm) of the *MoS*_2_ suspension two peaks were observed at 682nm and 715nm. The peak at 682 nm equals 1.83 eV, which corresponds the presence of a single layer of *MoS*2 and the peak at 715 nm equals 1.73 eV, which corresponds to the presence of a double layer of *MoS*2 in the sample (18). When the same synchronous scan was performed with the nanocomposite, there was an amplification of bacterial fluorescence peak at 703 nm that corresponded roughly with the double layer photoluminescence. Hence we may consider it to be a resonance peak. When the difference spectrum was calculated that is *δF* = bacteria-*MoS*_2_ – only bacteria, a peak was obtained which corresponded exactly with the 685 nm peak.

### Current voltage (I-V) characteristics

Graphene-biofilm samples exhibited current of the order of 10^−^4*A*. This corresponds well with reports that show increased conductivity of hydrogels when graphene is incorporated. Graphene-*MoS*_2_ hybrid have shown an exceptionally high current when incorporated into the photosynthetic biofilms, of the order of 10^−^3*A*. This was much greater than current

The threshold of graphene-*MoS*2-biofilm was 2.4 V. Biofilms incorporated with either graphene or *MoS*_2_ exhibited a threshold of 0.75 V and 0.85 V respectively. The threshold in case of graphene-biofilm was not very prominent since this is a highly defective graphene sample and graphene does not have a direct band gap material. The band gap in case of *MoS*_2_ graphene biofilm composite is more prominent because there are two Schottky junctions present: gold-*MoS*_2_ junction and the graphene-*MoS*_2_ junction. Control sample did not have any threshold value.

### Impedance Spectroscopy

In case of AC stufies impedence could be a combination of resistance, capacitance and inductance. We can reconstruct the equivalent circuit using the knowldge on real and imagianry part of impedance. The effect of bias voltages are seen in figure (5).

In figure (5) there seems to be a critical bias voltage beyond which the Nyquist diagram drastically changes. This bias (close to 3-4V) is closer to the voltage at which the gating occurs (see figure (4)).At 4V bias, 2 semicircles were observed with an absence of Warberg impedence element for both control and *MoS*2 sample (see figure 5, panel C). Graphene (see figure 5, panel B) on the other hand showed a systematic variation of the Warburg-less semicircle pattern, with the semiciclr radii shrinking.

**Figure 4:**
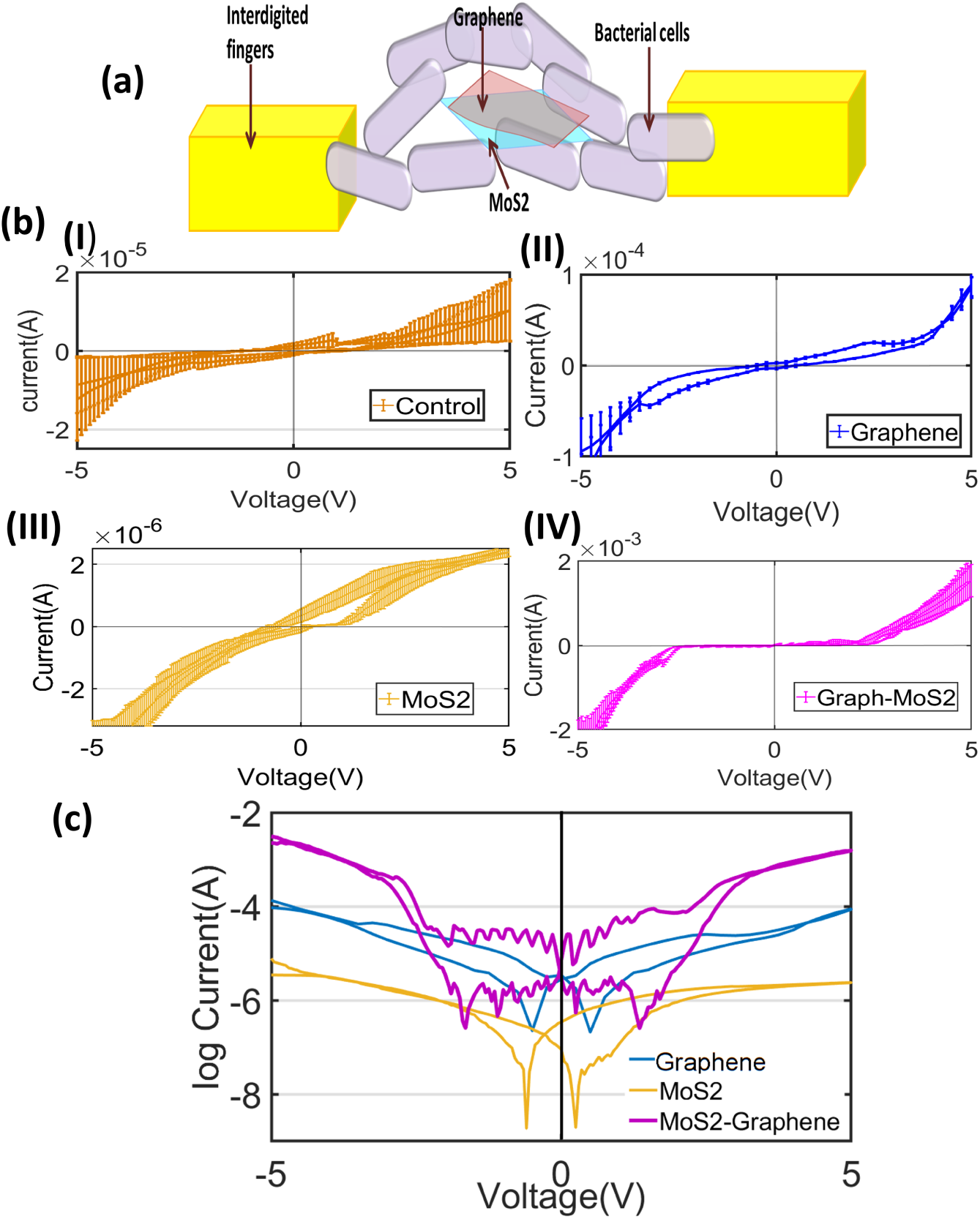
(a)Schematic showing bacterial cell and nanomaterials distribution between two IDE fingers (b) Current-voltage curves for (I) control,(II) graphene-biofilm, (III)*MoS*2-biofilm and (IV) *MoS*2+graphene biofilm hybrids, respectively (C) Current-voltage plots for nanomaterial incorporated samples in log scale for easy comparison of current values through any of the other samples. Such increase in the current of graphene-MoS2 sample was observed as compared to *MoS*_2_ or graphene due to additive effect of both *MoS*2 and graphene.

**Figure 5:**
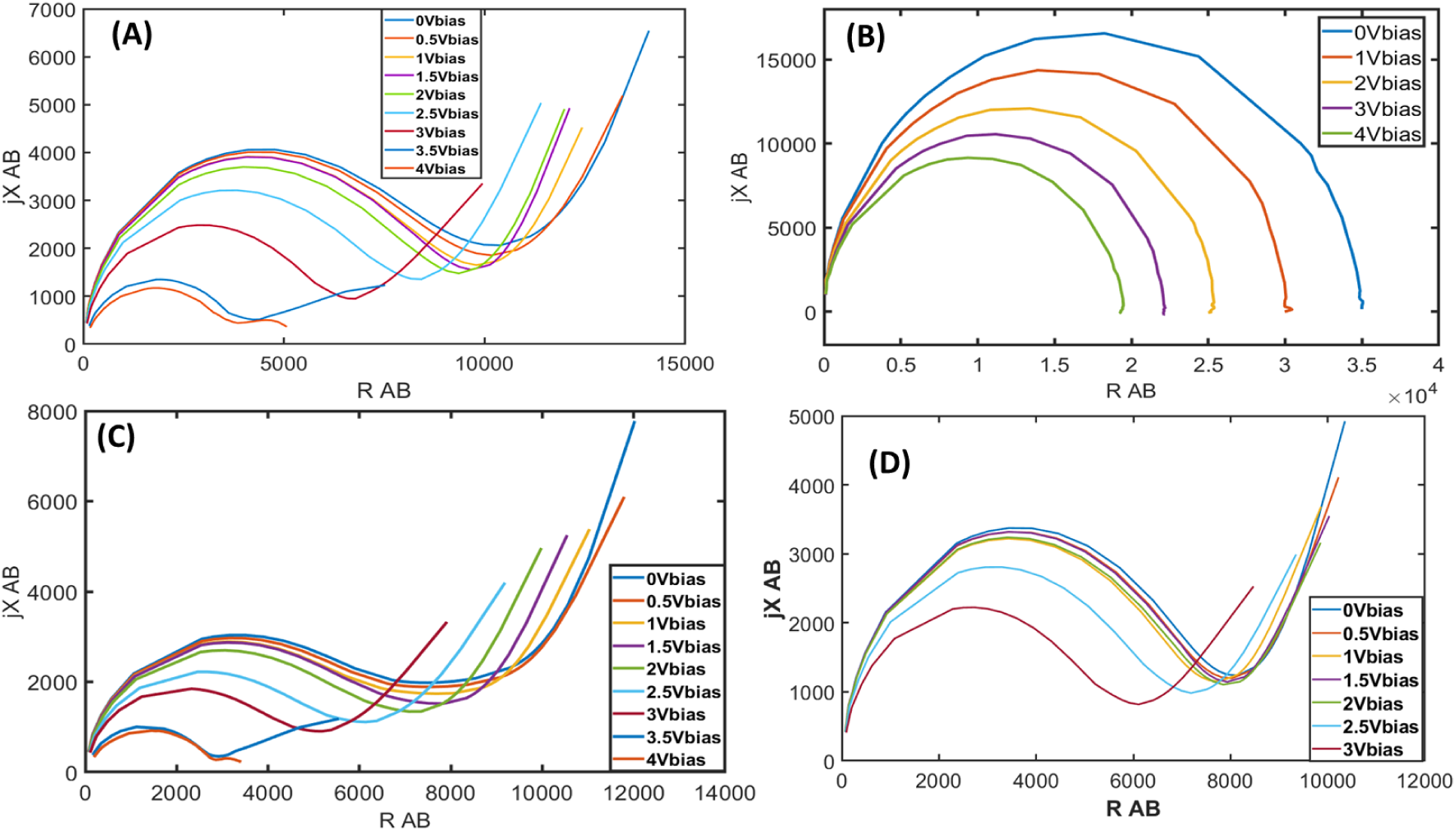
Nyquist plot of biofilm only(A), Graphene biofilm(B), *MoS*2 biofilm(C), *MoS*2+graphene biofilm(D) at different bias voltage.

**Figure 6:**
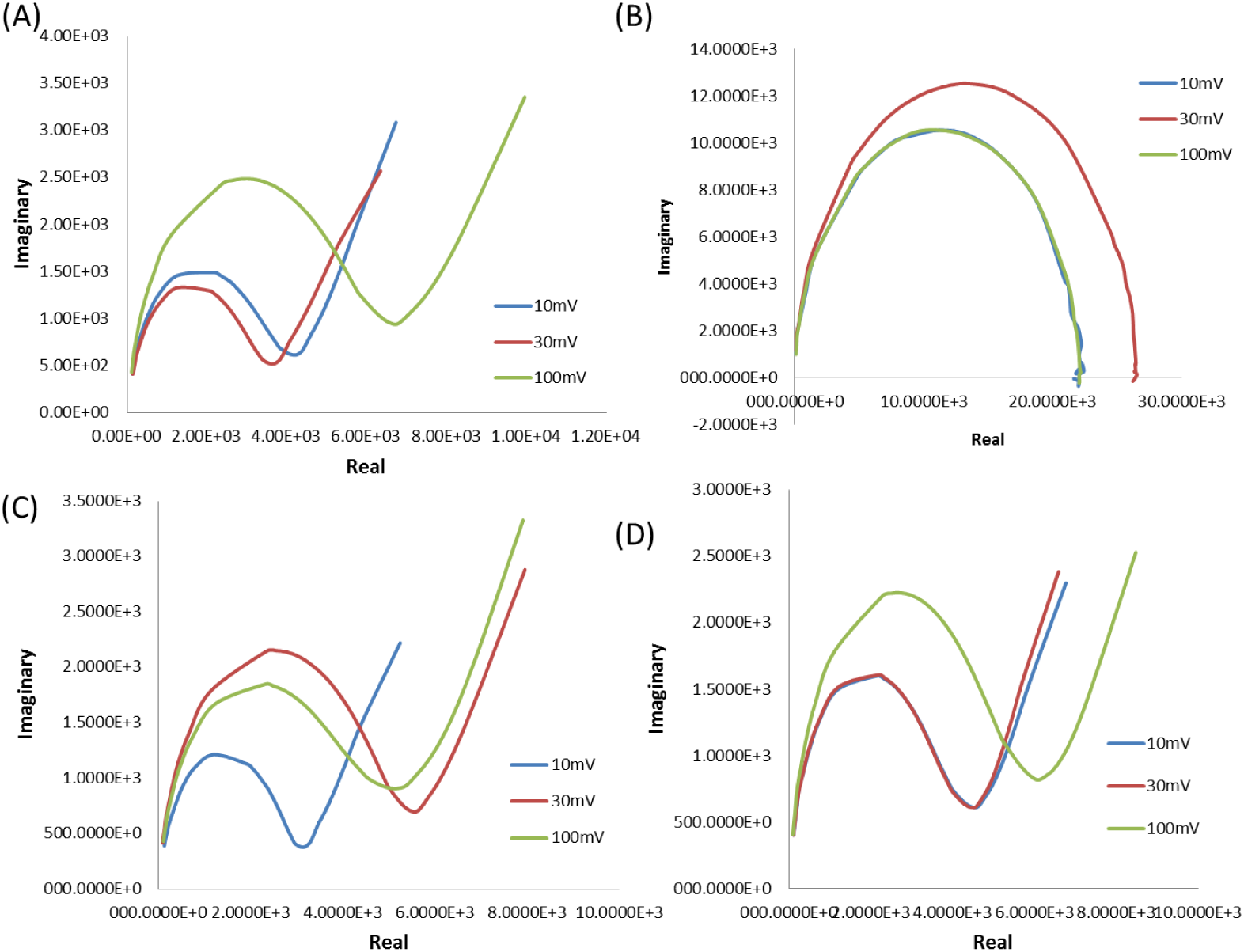
RMS voltage Variation at 10mV, 30mV and 100mV of biofilm only(A),Graphene biofilm(B),*MoS*2 biofilm(C) and *MoS*2-Graphene biofilm(D).

The variation of RMS voltage for a given bias (3V) may be noteworthy.

What follows is that the overall impedence behaviour drives the electrical properties of even the nanohybrid biofilms (20). When the RMS voltage is increased from 10mV to 30mV there is hardly any change in the control biofilm (see 6,panel A), whereas for *MoS*2 ((see 6,panel C)) there is a big jump of the nyquist plot. In contrast the 10 to 30mV jump is attenuated in M+G (see 6,panel D). The graphene only panel (see 6,panel B)) maintains uniqueness, namely absence of any Warburg element. The criticality in the RMS voltage is only reflected in slight change in the semicircle radis of the nyquist diagram.

Figure (7) represents a comparative appraisal of the Nyquist and Bode diagram for the respective semiconducting biofilms (see panels A & B of figure(7). The panel B suggests that Mos biofilm has a characteristic Warburg behavior, suggesting distinctive electric behavior of such semiconducting film.Eventually, graphene exhibited the maximum diameter of the semicircle with the Warburg impedence element absent. The Bode diagram (7,panels C & D) showed a distinctive frequency response for the graphene (flat for real part of the impedance, panel C, and very sharp for imaginary part of the impedance, panel D).

**Figure 7:**
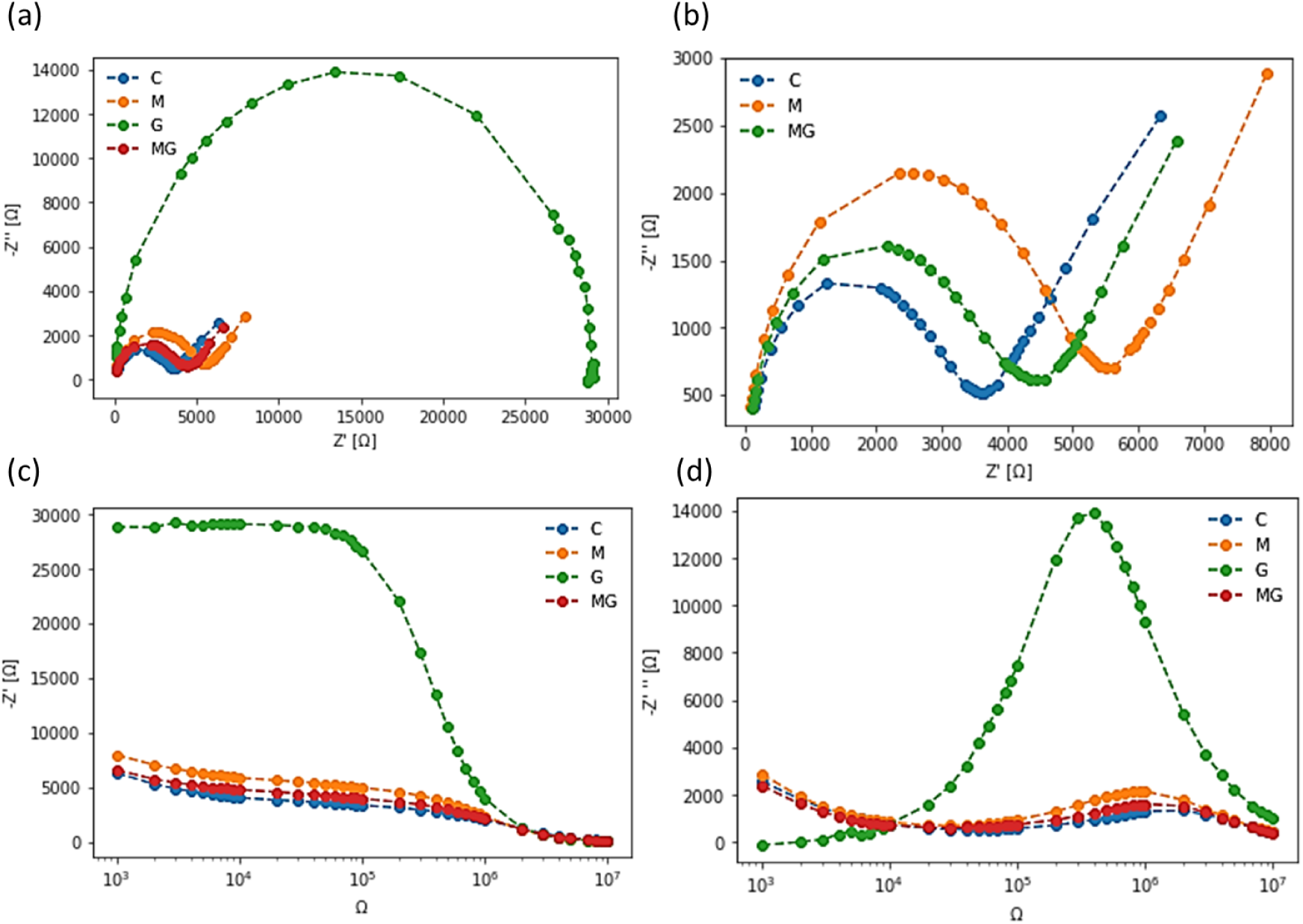
Nyquist plots real v/s imaginary at 3V DC bias. Biofilm only(C),*MoS*2 with Biofilm(M),Graphene with biofilm(G), *MoS*2+Graphene with biofilm(MG) (a). same without graphene biofilm(b). real (-z’)v/s frequency at 3V DC bias(c). imaginary(-z”) v/s frequency at 3V DC bias (d).

## Discussions

In this paper, commercially available, bulk *MoS*2 was subjected to liquid phase exfoliation in order to obtain a suspension of *MoS*2 nanosheets. The obtained material was characterized by various spectroscopic techniques and found to be constituted of a mixture of single and double-layered *MoS*2 nanosheets. However, the suspension of *MoS*2 was not very stable and tended to precipitate within a few days due to the re-stacking of the exfoliated layers. In order to obtain a more stable suspension, commercially available graphene nanoplatelets, known to form stable dispersions in water, were added to the *MoS*2 prior to sonication. Thus a suspension of graphene-*MoS*2 heterojunction materials was formed which showed very high stability even after a month. The Nyquist diagram also suggests that the nyquist pattern is more stable

### A semiconducting biofilm and its emergent properties

Previously (1) we have demonstrated the emergent semiconducting properties of graphene when immobilized within the adherent biofilm. However, graphene is an indirect bandgap material, and the emergence of the bandgap is dependent on a number of factors. No threshold was observed with the commonly available, cheaper, more defective forms of graphene as used in this paper. Therefore, we decided to work on other 2D material with well known direct bandgap and hence semiconducting and photoluminescent properties.

Electrical properties were measured for four kinds of samples with identical device dimensions. Samples were control biofilms, *MoS*2+biofilm, graphene+biofilm and *MoS*2+graphene+biofilm.

Graphene-biofilms showed higher conductivity than *MoS*2 biofilms since graphene itself is a good conductor. The effects of immobilization of graphene within the biofilm (and consequent strain on the carbon lattice) were apparent from the non-linearity in its conductivity at lower voltages. However, there was no apparent threshold in current conduction.

For *MoS*2-biofilms there was a clear threshold on one side of the curve and not the other side. This threshold indicates a Schottky junction like electrical interface formed between the metal electrode and the semiconducting *MoS*2 nanosheet. However, the threshold was small compared to what has been observed previously with graphene. A smaller threshold coupled with the fact that nanosheets are added as a dispersion (hence the equal probability of its immobilization on either side of the inter-digitated electrode) simply means that insufficient quantities of *MoS*2 were immobilized or that their dispersion was too un-uniform, as confirmed by visual inspection.

Hence, *MoS*2 and graphene (defective) were-co-exfoliated and the higher number of polar groups on the defective graphene helped to keep the hydrophobic *MoS*2 stable within an aqueous solution. When *MoS*2+graphene heterojunction complexes were immobilized within biofilms, the current-voltage plot showed a very distinct and symmetrical threshold. This signifies gating of current flow through a Schottky diode like an interface between the metal electrode *MoS*2 -> graphene. The precise symmetrical nature of the material signifies very good dispersion and subsequent insertion of *MoS*2 into the biofilm. This is the first demonstration of a live, photosynthetic biofilm showing semiconducting behavior.

The effect of *MoS*2 on the intrinsic fluorescence of biofilms merits some discussion. Supplementary figure (S2) confirms that there is some enhancement of fluorescence in the NIR region

## Conclusion

We have been successful in obtaining a semiconducting bacterial biofilm based on a direct bandgap nanomaterial namely *MoS*2. The emergence of threshold upon biofilm mediated immobilization of *MoS*2 on metal electrodes is a direct indicator of the semiconducting nature of the whole composite. Further our work demonstrates that biofilms have been remarkably efficient in supporting an embedded network of nanoscale Schottky diodes. Since the biofilm is of photosynthetic origin, the resultant material may be more efficient solar energy harvesting applications. The possibility of using the material as a wound healing patch, coating for important drugs and enzymes may also be of some future interest.

## Supporting information

Fig S1: Growth Curve Fig S2: Fluorescence Amplification

## Acknowledgement

I thank Dr. Achintya Singha of Bose Institute and Prof. Sanatan Chattopadhyay of the Electronics Department, Rajabazar Science College campus of the University of Calcutta. We also thank Prof. Sanjay Ghosh, the Head Department of Biochemistry and all the colleagues therein for their support.

