## Supplementary material for "Semiconducting bacterial biofilm based on graphene-MoS2 template and component dependent gating behavior": Fig S1: Growth Curve Fig S2: Fluorescence Amplification

### Additional Material for Semiconducting bacterial biofilm based on graphene-Mos2 template and component dependent gating voltage


Sanhita Ray,[†,‡] Arpita Das,[†] and Anjan Kr Dasgupta[*,†]

†*Department of Biochemistry, University of Calcutta, Kolkata*

‡*Department of Electrical Engineering, Jack Baskin School of Engineering, University of California, Santa Cruz*


**Abstract**

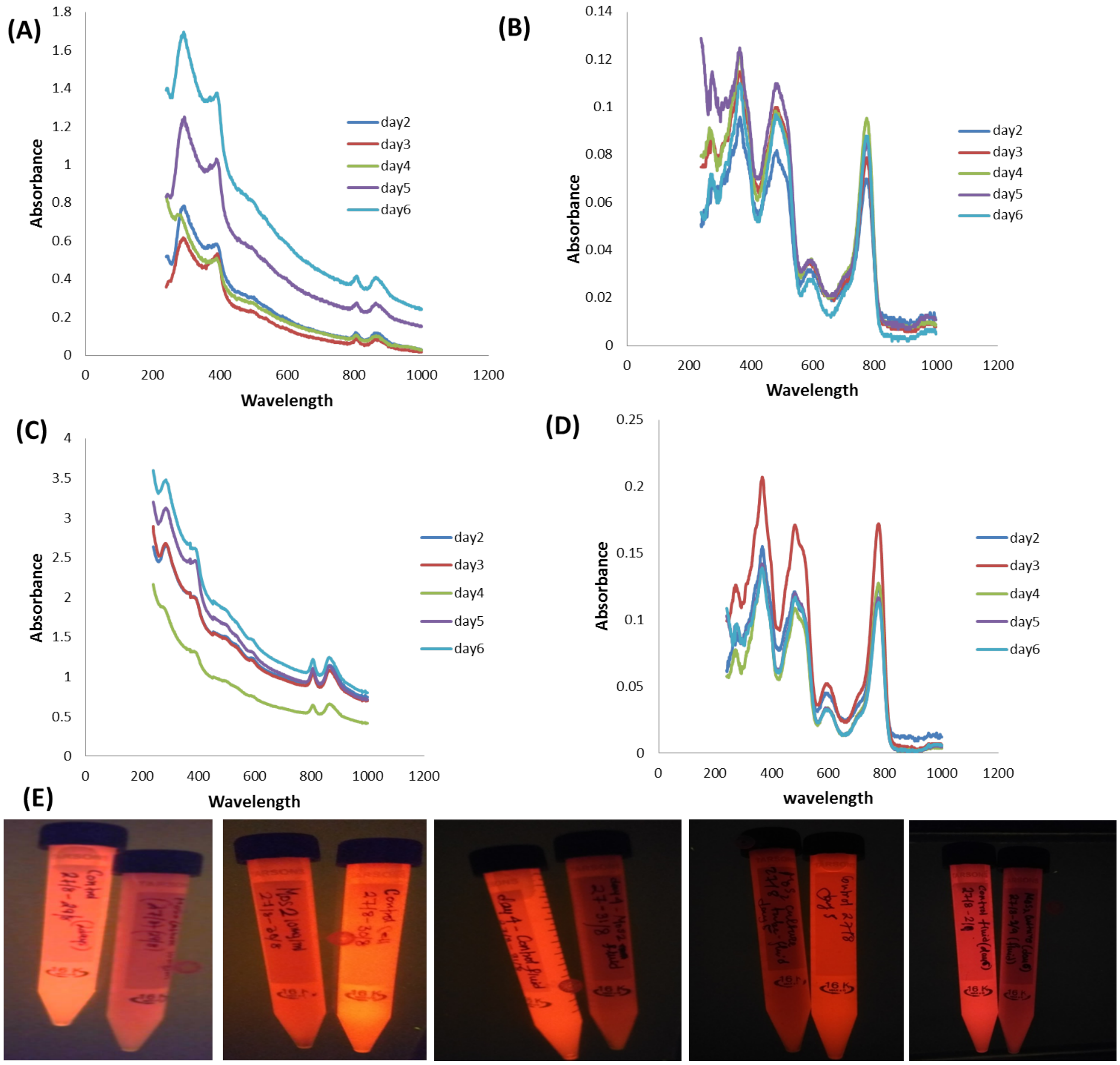


Figure 1: Fig S1: Growth Kinetics study. Daywise Absorbance spectra of planktonic cells (A) and pigments(B) of Biofilm. Daywise absorbance spectra of planktonic cells(C) and pigments(D) of biofilm in presence of MoS2. Daywise growth from day-2(left) to day-6(right) observed under UV.

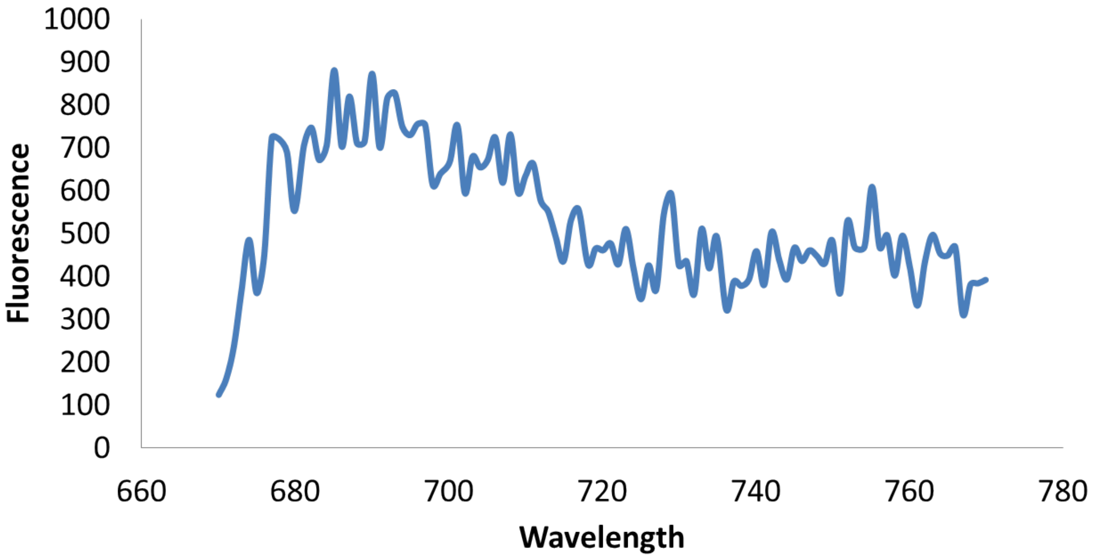


Figure 2: Fig S4: Fig S4: Synchronous scan (438-538:670-770nm) of MoS2 obtained from MoS2 biofilm by substracting MoS2-biofilm from biofilm.
